# Establishment of an African green monkey model for COVID-19

**DOI:** 10.1101/2020.05.17.100289

**Authors:** Courtney Woolsey, Viktoriya Borisevich, Abhishek N. Prasad, Krystle N. Agans, Daniel J. Deer, Natalie S. Dobias, John C. Heymann, Stephanie L. Foster, Corri B. Levine, Liana Medina, Kevin Melody, Joan B. Geisbert, Karla A. Fenton, Thomas W. Geisbert, Robert W. Cross

## Abstract

Severe acute respiratory syndrome coronavirus 2 (SARS-CoV-2) is responsible for an unprecedented global pandemic of COVID-19. Animal models are urgently needed to study the pathogenesis of COVID-19 and to screen candidate vaccines and treatments. Nonhuman primates (NHP) are considered the gold standard model for many infectious pathogens as they usually best reflect the human condition. Here, we show that African green monkeys support a high level of SARS-CoV-2 replication and develop pronounced respiratory disease that may be more substantial than reported for other NHP species including cynomolgus and rhesus macaques. In addition, SARS-CoV-2 was detected in mucosal samples of all animals including feces of several animals as late as 15 days after virus exposure. Importantly, we show that virus replication and respiratory disease can be produced in African green monkeys using a much lower and more natural dose of SARS-CoV-2 than has been employed in other NHP studies.

## Introduction

Severe acute respiratory syndrome coronavirus 2 (SARS-CoV-2), the etiological agent of coronavirus disease 2019 (COVID-19), emerged in Wuhan, China in late 2019 and rapidly swept the globe. As of May 16, 2020, over 4.4 million confirmed cases and 302,000 deaths have been reported worldwide [1]. The COVID-19 crisis has inflicted an immense toll on human health causing profound socioeconomic effects that will linger for years to come.

Vaccines and treatments against SARS-CoV-2 could drastically reduce COVID-19 transmission, saving lives and brightening prospects for economic recovery. No licensed countermeasures currently exist, although a number of clinical trials are underway. While clinical testing is a good predictor of preventative and drug efficacy, caution is warranted with this approach due to the potential for disease enhancement. Vaccine candidates against SARS-CoV-1, including inactivated whole-virus, recombinant DNA subunit, virus-like particle, and live virus vector-based vaccines, induced an immunopathology following immunization and experimental infection of mice, ferrets, and nonhuman primates (NHPs) suggesting hypersensitivity to virus components [2–5]. Given the strong genetic similarity of SARS-CoV-2 and SARS-CoV-1 [6], an appropriate surrogate for human COVID-19 is needed to safely evaluate prophylactics. A careful assessment in animal models could reveal possible immune complications elicited by vaccines and therapies prior to their release to the public. Moreover, animal models could help shed light on important aspects of the disease in ways that are not easily addressed or feasible in humans, such as how the virus spreads and interacts with the host immune system.

An ideal COVID-19 animal model would replicate all facets of human disease. Several animal species including mice, hamsters, ferrets, and NHPs were found to support SARS-CoV-2 replication and displayed varying degrees of non-lethal illness when the virus was delivered into the respiratory tract of these animals [7–13]. While each of these models has utility in the study of COVID-19, NHPs have the closest physiological resemblance to humans allowing a better comparison of host responses to infection. This genetic similarity has also contributed to the increased availability of reagents to perform in-depth analyses of the immune response. Recently, the first studies evaluating the pathogenic potential of SARS-CoV-2 in cynomolgus and rhesus macaques were performed. Rhesus macaques developed pneumonia and clinical signs whereas disease in cynomolgus macaques was fairly mild indicating the former appears to better reflect more severe cases of COVID-19 [10–12]. These results suggest certain NHP species may serve as better models than others for coronavirus infections. For SARS, the disease caused by SARS-CoV-1, African green monkeys (AGMs) were found to support the highest level of viral replication, followed by cynomolgus macaques and rhesus macaques when all three species were challenged in parallel [14]. Only AGMs had notable replication in the lower respiratory tract following SARS-CoV-1 inoculation; necropsy of these animals indicated focal interstitial mononuclear inflammatory infiltrates and edema in the lung consistent with human SARS. As SARS-CoV-1 and SARS-CoV-2 share the same putative host receptor angiotensin-converting enzyme 2 (ACE2) [15, 16], we reasoned that AGMs might serve as a useful model for COVID-19.

Here, we infected AGMs with a low passage isolate of SARS-CoV-2 (SARS-CoV-2/INMI1-Isolate/2020/Italy) and evaluated their potential as a model for COVID-19. SARS-CoV-2/INMI1-Isolate/2020/Italy was isolated from the first clinical case in Italy [17] and is the first V clade virus (GISAID) to be experimentally inoculated into NHPs. We demonstrate AGMs mimic several aspects of human disease including a high degree of viral replication and severe pulmonary lesions. This model can be used to conduct pathogenesis studies and to screen potential vaccines and therapeutics.

## Results

We challenged six adult AGMs with 5.0 x 10^5^ PFU of the Italy isolate of SARS-CoV-2 (SARS-CoV-2/INMI1-Isolate/2020/Italy) by combined intratracheal (i.t.) and intranasal (i.n.) routes (dose divided equally). A cohort of three animals was euthanized at 5 dpi, while the remaining three animals were held indefinitely. Blood from all animals was sampled on days 0, 2, 3, 4, and 5, and continuing with days 7, 9, 12, 15, and 21 for animals held past 5 dpi. No overt clinical signs of disease were observable in any of the animals, other than decreased appetite compared to baseline (0 days post-infection; dpi) in 5/6 animals, and a brief period of elevated body temperature suggestive of fever in two of three animals at 3 dpi (Supplementary Figure 1). A biphasic increase in partial O^2^ pressures were noted in two animals over the course of the study, but no overt changes in partial CO^2^ pressures in any animals were noted (Figure 1-A,B). Transient shifts in leukocyte populations and mild thrombocytopenia were observed in all animals, most prominently during days 2-7 post-infection (Supplementary Table 1, Figure 1-C,D,E,F). Markers for renal and hepatic function mostly remained unchanged (< 2-fold increases in ALT, AST, ALP); however, CRP, a marker of acute systemic inflammation, was elevated two to seven-fold in all animals 2-5 dpi (Table 1, Figure 1-G,H,I,J). Thoracic radiographs were taken on −1, 2, 3, 4, 5 dpi, and were shown to be inconclusive (Supplementary Figure 2). Gross examination at necropsy of three AGMs at 5 dpi revealed varying degrees of pulmonary consolidation with hyperemia in all monkeys (Figure 2). Lesions were multifocal to locally extensive within each lobe. The most severe lesions were located in the dorsal aspects of the lower lung lobes in all animals. A board-certified veterinary pathologist approximated lesion severity for each lung lobe (Supplementary Table 2) Small intestines in all monkeys were segmentally flaccid and distended with gas and yellow liquid contents. Mild lymphoid enlargement was noted in one of the three monkeys. There were no other significant gross lesions.

**Figure 1.**
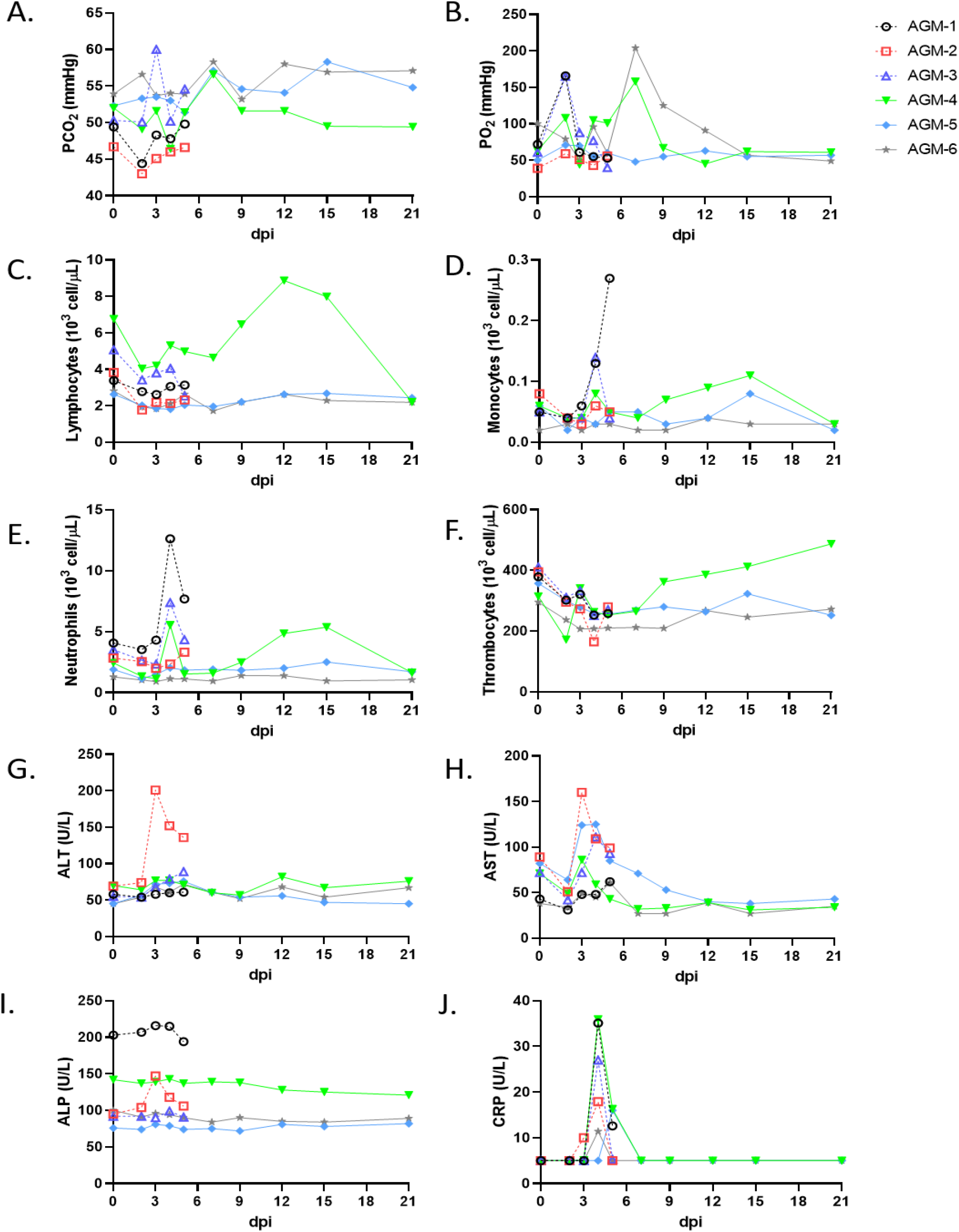
Hematological features of AGMs infected with SARS-CoV-2. Blood was collected from AGMs on procedure days (0, 2, 3, 4, 5, 7, 9, 12, 15, 21). Partial pressures of CO_2_ and O_2_ were obtained using an iSTAT Alinity hematological analyzer (Abbott). Leukocyte subpopulations and thrombocytes were differentiated and quantified using a Vetscan HM5 hematological analyzer (Abaxis). Serum analytes (ALT, AST, ALP, CRP) were quantified using a Piccolo serum biochemistry analyzer (Abaxis). For all graphs, dashed lines and open symbols (AGM-1, AGM-2, AGM-3) indicate AGMs sacrificed 5 dpi; solid lines (AGM-4, AGM-5, AGM-6) indicate AGMs held to 21 dpi.

**Figure 2.**
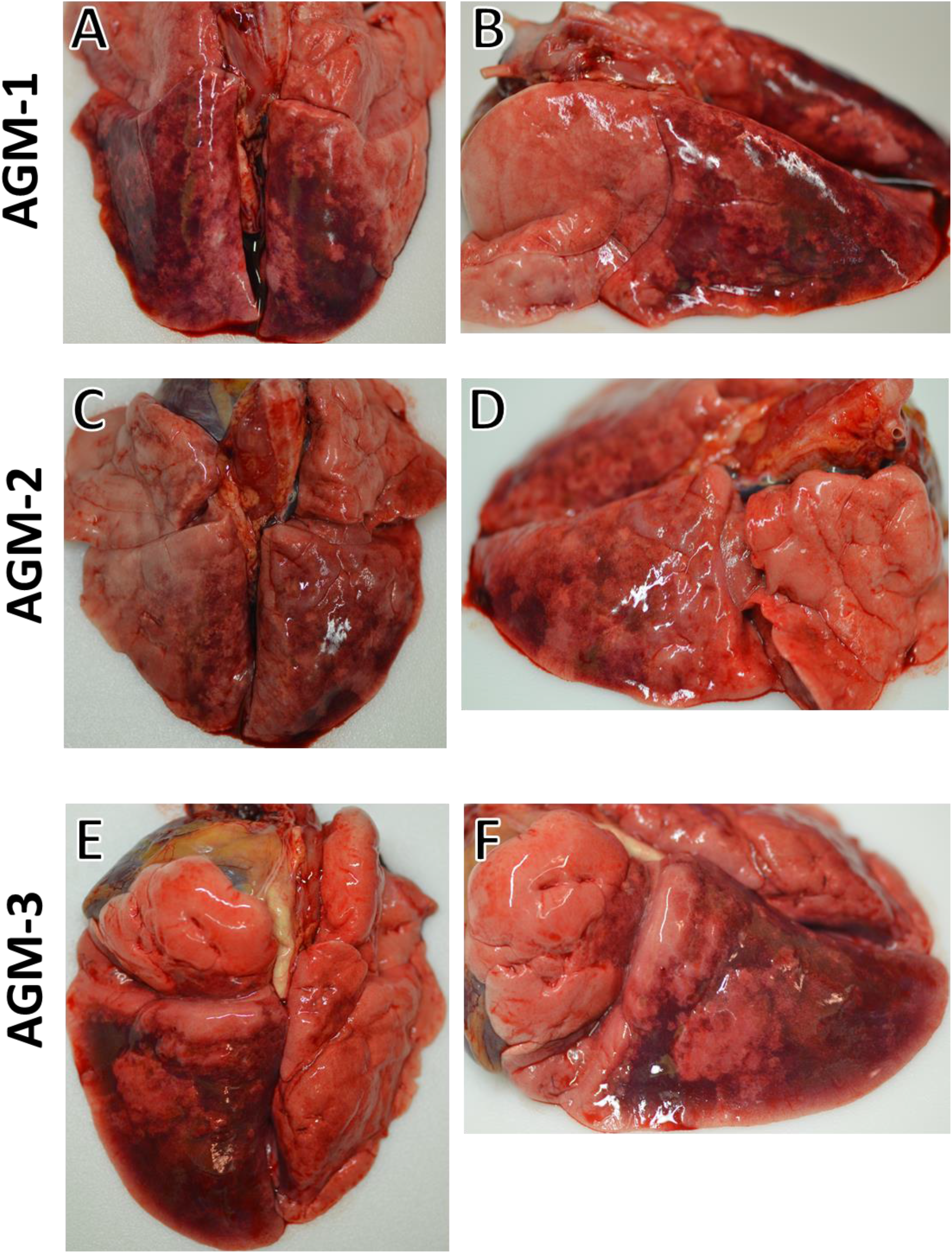
Lung gross pathology in SARS-CoV-2-infected AGMs. Side and dorsal views of lungs from AGMs sacrificed 5 dpi after infection with SARS-CoV-2 exhibiting marked pulmonary consolidation with hemorrhage, ranging from multifocal to locally extensive in severity. **(A)** and **(B)** are dorsal and side views of the lungs from AGM-1. **(C)** and **(D)** are dorsal and side views of the lungs from AGM-2. **(E)** and **(F)** are dorsal and side views of the lungs from AGM-3.

Histologically, all three AGMs at 5 dpi developed varying degrees of multifocal pulmonary lesions. In the most severely affected monkey, histologic features of pneumonia include acute inflammation centered on terminal bronchioles and numerous multinucleated giant cells with irregularly distributed nuclei. Continuous with the terminal bronchiolitis was evidence of diffuse alveolar damage (DAD) with hyaline membrane formation, type II pneumocyte hyperplasia, pulmonary edema and pulmonary hemorrhage. Occasionally, small fibrous tissue proliferations were noted within the terminal airways reminiscent of bronchiolitis obliterans organizing pneumonia (BOOP)-like lesions. Interstitial pneumonia was noted in the lesser affected regions of the lung along with congestion and increased numbers of alveolar macrophages. In the less affected monkeys, pulmonary lesions lack acute inflammation within the terminal bronchioles; however, interstitial pneumonia is present and in one monkey, multinucleated giant cells with irregularly distributed nuclei were prominently displayed within bronchioles. Colocalization of SARS antigen with pulmonary lesions were demonstrated with antibodies against SARS N protein. Positive immunohistochemical labeling was noted diffusely within the cytoplasm of alveolar macrophages, respiratory epithelium, type I pneumocytes and type II pneumocytes. One animal had only positive immunolabeling in respiratory epithelium. Genomic SARS-CoV2 RNA was detected by i*n situ* hybridization in pneumocytes and associated with the alveolar macrophages within acute inflammation centered on terminal bronchioles (Figure 3).

**Figure 3.**
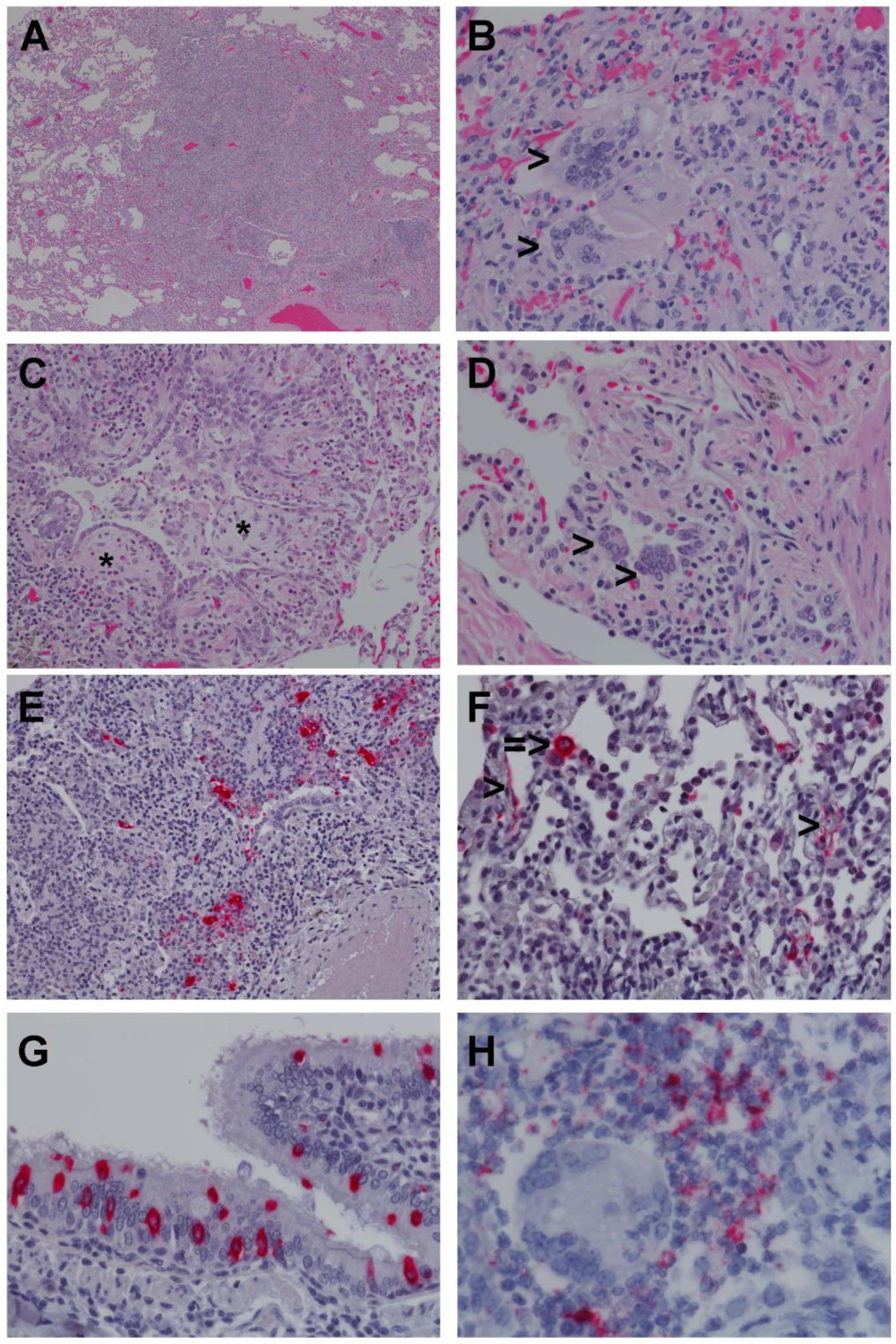
Pulmonary histologic changes in AGMs infected with SARS-CoV2. (A) Low magnification image displaying the marked acute bronchiolitis and interstitial pneumonia (4x) (B) Higher magnification image displaying multiple multinucleated giant cells (>) and acute bronchiolitis with edema and hemorrhage (40x) (C) Terminal bronchioles with multiple fibrous tissue proliferations reminiscent of bronchiolitis obliterans organizing pneumonia (BOOP)-like lesions (*) (20x) (D) Multiple multinucleated giants cells (>) within terminal bronchioles of lesser affected monkey (40x) (E) SARS-CoV 2 positive immunohistochemical (IHC) labeling (red) of alveolar macrophages and pneumoncytes within acute bronchiolar inflammation (20x) (F) SARS-CoV 2 positive IHC labeling (red) of type I (>) and type II (=>) pneumocytes (40x) (G) SARS-CoV 2 positive IHC labeling (red) of respiratory epithelium (40x), (H) Genomic SARS-CoV2 RNA (red) detected by *in situ* hybridization in a cluster of cells associated with acute bronchiolitis (60x).

We quantified viral load from mucosal swabs and bronchoalveolar lavage (BAL) fluid collected on procedure days by RT-qPCR and plaque titration (Figure 4). All animals had detectable quantities of vRNA and infectious virus from nasal secretions beginning at 2 dpi, with viral titers ranging from ~2-4 log PFU/mL (Figure 4-A,B). Both viral RNA (vRNA) and infectious virus were detected from oral swabs from only three animals (AGM-2, AGM-5, AGM-6) 3 dpi, and persisted in AGM-5 up to 7 dpi (Figure 4-C,D). Infectious virus was only detectable from the rectal swab of a single animal (AGM-3) 3-5 dpi; however, vRNA was also detected from AGM-4 and AGM-5 (Figure 4-E,F). Both vRNA and infectious virus was present in BAL fluid from all animals beginning at 3 dpi (Figure 4-G,H). Neither vRNA nor infectious virus was detected by RT-qPCR of RNA from whole blood or plaque titration of plasma from any animal, indicating a lack of circulating cell-associated or free virus in the peripheral blood (data not shown).

**Figure 4.**
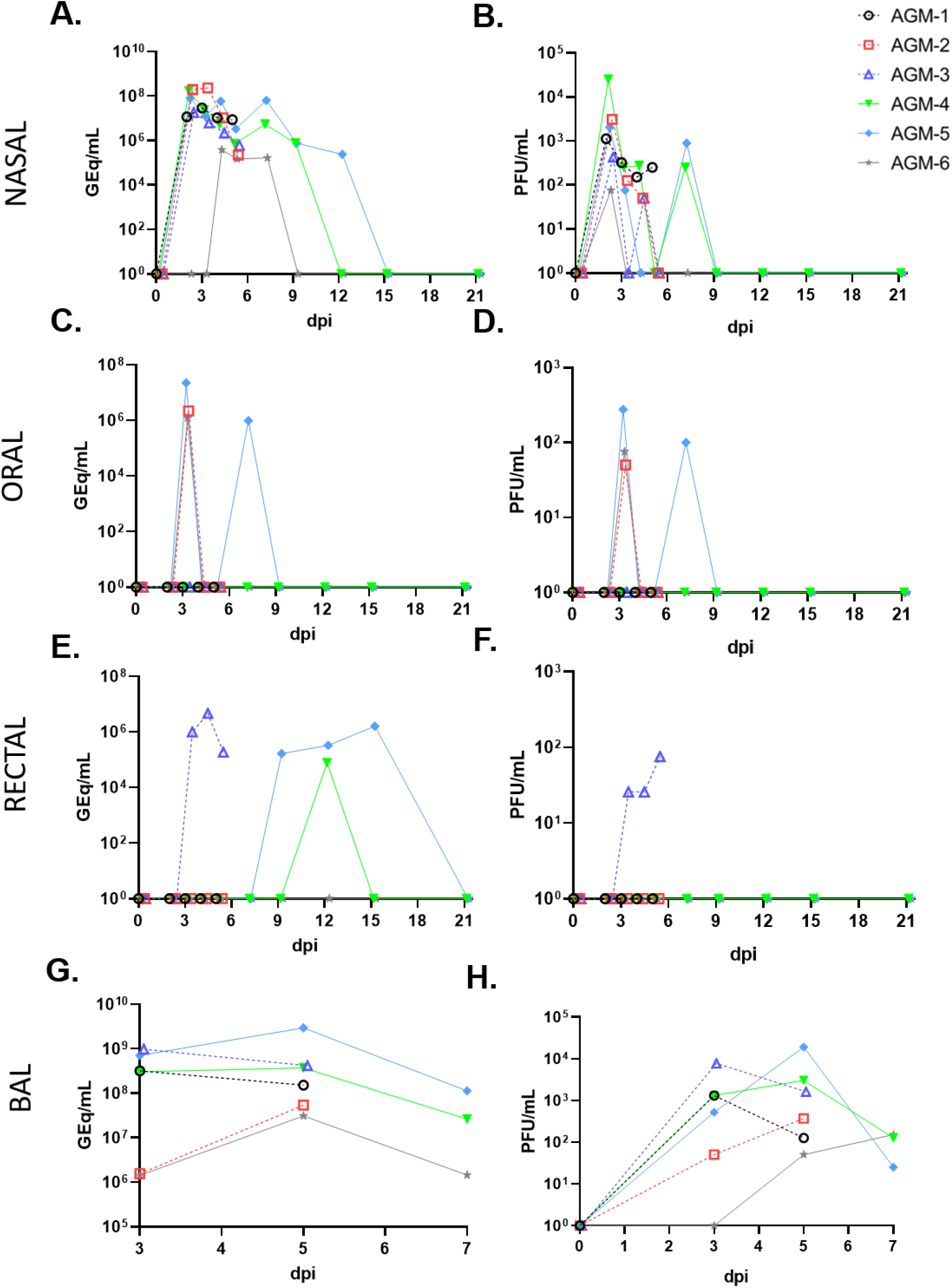
Detection of SARS-CoV-2 vRNA and infectious virus in muscosal swabs and BAL fluid. RNA extracted from swabs of mucosa and BAL fluid from AGMs infected with SARS-CoV-2 was subjected to RT-qPCR analysis or viral titration. **(A,B)** nasal swabs, **(C,D)** oral swabs, **(E,F)**rectal swabs, **(G,H)** BAL fluid. For all graphs, dashed lines and open symbols (AGM-1, AGM-2, AGM-3) indicate AGMs sacrificed 5 dpi; solid lines (AGM-4, AGM-5, AGM-6) indicate AGMs held to 21 dpi. Data plotted is the mean of duplicate RT-qPCR reactions or duplicate wells.

Similar to human cases of COVID-19, systemic concentrations of a number of inflammatory cytokines and chemokines were elevated following AGM infection. Serum levels of pro-inflammatory IL-8, IP-10, IL-12, and monocyte chemoattractant protein (MCP-1) peaked at 2 DPI corresponding to subsequent recruitment of monocytes and neutrophils in the blood (Figure 1-D,E, Figure 5-C,D,E,F). Increased secretion of IL-6, IFN-beta, and IL-10was also evident at this time point (Figure 5-A,G,H). Notably, IL-6 is thought to predict respiratory failure in human COVID-19 cases and may serve as an effective immunological biomarker [18]. As IL-6 is a main regulator of acute phase fibrinogen synthesis, and elevated fibrinogen and other coagulation abnormalities are thought to correlate with disease severity in hospitalized patients [19, 20], we also measured levels of this clotting factor. Fibrinogen levels surged in 4 of 6 monkeys indicating potential coagulation abnormalities in these animals (Figure 5-B). This increase in circulating fibrinogen aligns with our gross pathology findings of substantial hemorrhage in the lung of monkeys euthanized at 5 DPI. Collectively, these results indicate the host response to SARS-CoV-2 infection in AGMs parallels that of humans.

**Figure 5.**
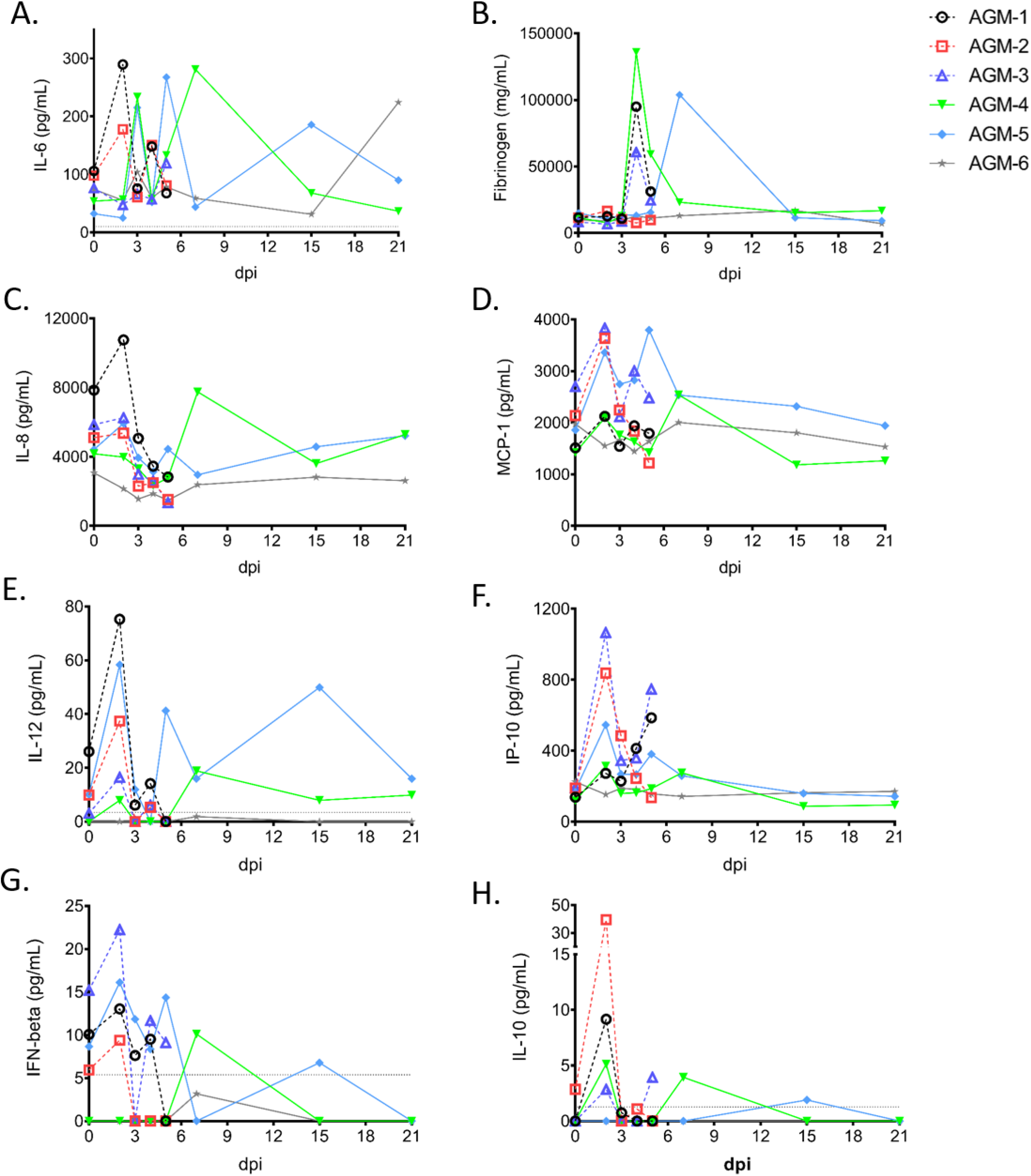
Soluble markers of inflammation detected in SARS-CoV-2 infected AGM sera over the course of the study.

Four animals seroconverted against SARS-CoV-2, including all three animals held past 5 dpi, with the earliest detection of anti-SARS-CoV-2 IgG occurring at 5 dpi in AGM-3, AGM-4, and AGM-6 (Figure 6-A). IgG titers peaked at 15 dpi, with AGM-6 exhibiting low titers at 15 and 21 dpi compared to AGM-4 and AGM-5. Neutralizing titers (50% plaque reduction values) ranged from 1:8-1:32 were evident beginning at day 5 in one animal and all three animals followed to the study endpoint (Figure 6, B,C,D,E).

**Figure 6.**
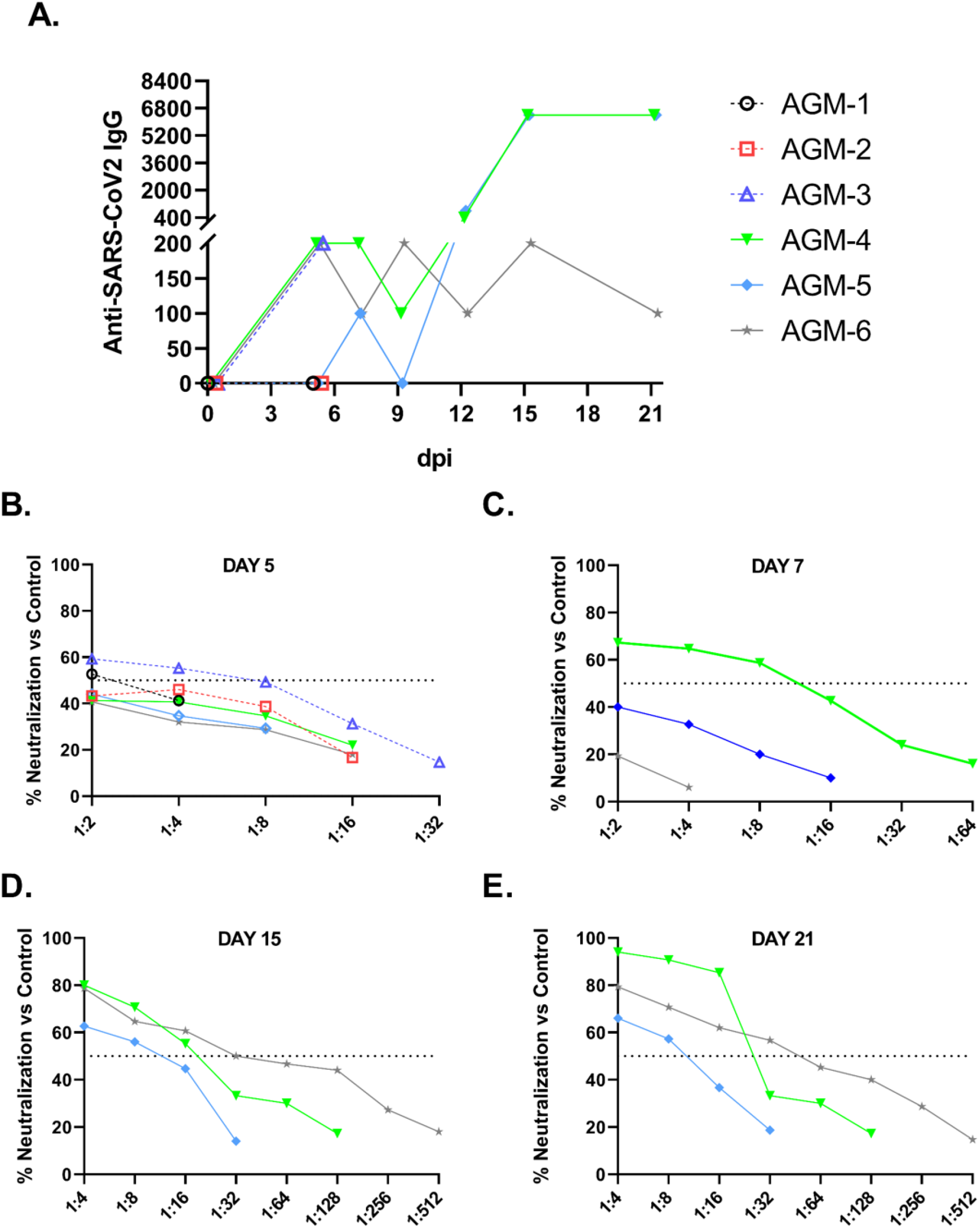
Serum anti-SARS-CoV-2 antibody titers and virus neutralization in AGMs. Anti-SARS-CoV-2 IgG binding titers were determined using traditional ELISA methodology where the antigen was whole infected cell lysate (background subtraction was performed using virus negative cell lysate) and titers are represented as reciprocal dilutions (**A**.). Antibody neutralization titers were also determined for sera collected 5 dpi **(B)**, 7 dpi **(C)**, 15 dpi **(D)**, and 21 dpi **(E)**. Data plotted represents the percent reduction in plaque counts (average of duplicate wells) compared to the average of three independent virus control plates (average plaque count across all three plates = 75). Dashed lines on each graph indicate 50% neutralization compared to the virus control plates. All plaque counts were calculated from duplicate wells at each dilution.

## Discussion

The emergence of SARS-CoV-2, and its astonishingly rapid evolution from localized outbreak to pandemic threat has emphasized the importance of coronaviruses with regards to human health globally, particularly of those clustering with lineage B betacoronaviruses. Accordingly, a necessity exists for the development of animal models for SARS-CoV-2 which recapitulate the COVID-19 disease phenotype observed in humans. With regard to NHPs, efforts to do so have been made using both rhesus and cynomolgus macaques [11–13, 21], with rhesus macaques more closely exhibiting the symptoms of COVID-19 disease in humans. Given that AGMs have shown utility as disease models for a number of respiratory pathogens (i.e., Nipah [22], SARS-CoV-1 [14]), we investigated their suitability as a model for SARS-CoV-2 infection.

AGMs challenged with an isolate of SARS-CoV-2 from Italy did not develop overt, debilitating clinical illness; however, changes in leukocyte subpopulations, thrombocytes, and serum markers of acute inflammatory processes indicated a systemic response to infection. Using surgically implanted temperature data loggers we were able to detect elevated body temperatures in two animals 3 dpi, indicative of fever. Moreover, all animals held past 5 dpi seroconverted, with two animals achieving high IgG titers (1:6400). The third animal held to 21 dpi never had an anti-SARS-CoV-2 IgG titer above 1:200, suggesting a dampened humoral response in this animal, and that immune arms other than humoral immunity may be important during SARS-CoV-2 infection. Additionally, the potential for re-infection in humans with SARS-CoV-2 has been speculated, but the risk factors or actual incidence are unknown. However, antibody titers to endemic human coronaviruses (e.g., HCoV-229E) are reported to gradually wane and re-infection with the homologous virus has been reported [23, 24].

An important aspect of the AGM model for SARS-CoV-2 infection is the development of pronounced viral pneumonia. All AGMs in our study exhibited pulmonary consolidation with hemorrhage, varying in severity between animals and lung lobes. We performed thoracic radiographs on all six SARS-CoV-2-infected AGMs; however, the radiographs while consistent with work on SARS-CoV-2-infected macaques recently published by others [11, 12, 21] did not reflect the degree of lesions and hemorrhage of the lungs seen at necropsy. It is our opinion that the radiographs from our study as well as those from the other COVID-19 NHP studies do not convincingly demonstrate SARS-CoV-2-induced disease. Specifically, opacities and changes observed in radiographs could be nonspecific and due to atelectasis, poor inspiratory effort, and BALS collection procedures. Important to this discussion, the current recommendation in humans is to not even use radiography for the primary diagnosis of COVID-19. Notably, a recent study of 636 ambulatory patients with COVID-19 did not detect any abnormalities in chest X rays of almost 60% of the cases [25]. The use of computed tomography (CT) scanning or even plesmography techniques may be more informative for detailed pulmonary investigations using animal models.

There was evidence of gastrointestinal involvement in the SARS-CoV-2-infected AGMs, which has been reported in human cases [23], as the putative SAR-CoV-2 entry receptor ACE-2 is expressed in the epithelium of the ileum and colon [26]. However, while all animals in our study exhibited abnormalities within the small intestine, we did not observe signs of gastrointestinal distress during the study course, with the possible exception of decreased appetite in most animals.

Our data show that infection of AGMs with SARS-CoV-2 results in the release of inflammatory mediators with similar immune signatures as human cases. Several groups reported that levels of IL-6, IL-8, and IL-10 correlate with disease severity in human patients [27, 28]. In this study, we observed increased serum levels of these interleukins as well as other pro-inflammatory cytokines and chemokines elevated in human cases. Moreover, we detected a rise in fibrinogen in a majority of monkeys, another prominent finding in human COVID-10 cases. Fibrinogen is implicated in thrombosis and vascular injury suggesting this discovery warrants further investigation. Together, these results indicate the AGM model can be used to study the host response to COVID-19. The heterologous response of AGMs along with the ability to collect tissues and longitudinal samples permits a detailed study of pathogenesis and immunity to COVID-19.

## Methods

### Virus

The virus (SARS-CoV-2/INMI1-Isolate/2020/Italy) employed was isolated on January 30, 2020 from the sputum of the first clinical case in Italy, a tourist visiting from the Hubei province of China that developed respiratory illness while traveling [17]. The virus was initially passaged twice (P2) on Vero E6 cells; the supernatant and cell lysate were collected and clarified following a freeze/thaw cycle. This isolate is certified Mycoplasma and FMDV free. The complete sequence was submitted to GenBank (MT066156) and is available on the GISAID website (BetaCoV/Italy/INMI1-isl/2020: EPI_ISL_410545) upon registration. For *in vivo* challenge, the P2 virus was propagated on Vero E6 cells and the supernatant was collected and clarified by centrifugation making the virus used in this study a P3 stock.

### Animal challenge

Animals were anesthetized with ketamine and inoculated with ~5 × 10^5^ PFU of SARS-CoV-2 (SARS-CoV-2/INMI1-Isolate/2020/Italy) with the dose being equally divided between the intratracheal (i.t.) (5.0 ml) and the intranasal (i.n.) (1.0 ml total at 0.5 ml per nostril) routes for each animal. Animals were longitudinally monitored for clinical signs of illness including temperature (measured by surgically implanted DST micro-T small implantable thermo loggers (Star-Oddi, Gardabaer, Iceland), respiration quality, and clinical pathology. All measurements requiring physical manipulation of the animals were performed under sedation by ketamine. All animal studies were approved by the University of Texas Medical Branch (UTMB) Institutional Animal Care and Use Committee and adhere to the NIH Guide for the Care and Use of Laboratory Animals.

### Radiographic technique

All monkeys were imaged with a portable GE AMX-4+ computed radiography system using a DRTECH detector set at a 36 inch focal film distance. Images were captured and evaluated using the Maven Patient Image Software in ventral dorsal (VD) and right lateral (R LAT) positions at 50 kVp and 12.5mA as previously described [22]. Chest radiographs were captured and interpreted by a double board-certified clinical veterinarian and veterinary pathologist and reviewed by a MD board-certified radiologist.

### RNA isolation from SARS-CoV-2-infected AGMs

On specified procedure days (days 0, 2, 3, 4, 5, 7, 12, 15, 21), 100 μl of blood was added to 600 μl of AVL viral lysis buffer (Qiagen) for virus inactivation and RNA extraction. Following removal from the high containment laboratory, RNA was isolated from blood and swabs using the QIAamp viral RNA kit (Qiagen).

### Detection of SARS-CoV-2 load

RNA was isolated from blood and mucosal swabs and assessed using the CDC SARS-CoV-2 N2 assay primers/probe for reverse transcriptase quantitative PCR (RT-qPCR)[29]. SARS-CoV-2 RNA was detected using One-step probe RT-qPCR kits (Qiagen) run on the CFX96 detection system (Bio-Rad), with the following cycle conditions: 50°C for 10 minutes, 95°C for 10 seconds, and 45 cycles of 95°C for 10 seconds and 55°C for 30 seconds. Threshold cycle (*C_T_*) values representing SARS-CoV-2 genomes were analyzed with CFX Manager Software, and data are presented as GEq. To generate the GEq standard curve, RNA from Vero E6 cells infected with SARS-CoV-2/INMI1-Isolate/2020/Italy was extracted and the number of genomes was calculated using Avogadro’s number and the molecular weight of the SARS-CoV-2 genome.

Virus titration was performed by plaque assay with Vero E6 cells (ATCC CRL-1586) from all blood plasma and mucosal swabs, and BAL samples. Briefly, increasing 10-fold dilutions of the samples were adsorbed to Vero E6 cell monolayers in duplicate wells (200 μl). Cells were overlaid with EMEM agar medium plus 1.25% Avicel, incubated for 2 days, and plaques were counted after staining with 1% crystal violet in formalin. The limit of detection for this assay is 25 PFU/ml.

### Hematology and serum biochemistry

Total white blood cell counts, white blood cell differentials, red blood cell counts, platelet counts, hematocrit values, total hemoglobin concentrations, mean cell volumes, mean corpuscular volumes, and mean corpuscular hemoglobin concentrations were analyzed from blood collected in tubes containing EDTA using a Vetscan HM5 hematologic analyzer (Abaxis). Serum samples were tested for concentrations of albumin, amylase, alanine aminotransferase (ALT), aspartate aminotransferase (AST), alkaline phosphatase (ALP), blood urea nitrogen (BUN), calcium, creatinine (CRE), C-reactive protein (CRP), gamma-glutamyltransferase (GGT), glucose, total protein, and uric acid by using a Piccolo point-of-care analyzer and Biochemistry Panel Plus analyzer discs (Abaxis). Partial pressures of CO_2_ and O_2_ were obtained using an iSTAT Alinity hematological analyzer (Abbott).

### ELISA

Sera collected at the indicated time points were tested for SARS-CoV-2-specific immunoglobulin G (IgG) antibodies by ELISA. MaxiSorp clear flat-bottom 96-well plates (44204 ThermoFisher, Rochester, NY) were coated overnight with a 1:1000 dilution of irradiated SARS-CoV-2 infected or normal Vero E6 lysate in PBS, pH 7.4 kindly provided by Dr. Thomas W. Ksiazek (UTMB). Sera were initially diluted 1:100 and then four-fold through 1:6400 in 3% BSA in 1× PBS. After a one-hour incubation, plates were washed six times with wash buffer (1 x PBS with 0.2% Tween-20) and incubated for an hour with a 1:2500 dilution of horseradish peroxidase (HRP)-conjugated anti-primate IgG antibody (Fitzgerald Industries International, Acton, MA). RT SigmaFast O-phenylenediamine (OPD) substrate (P9187, Sigma, St. Louis, MO) was added to the wells after six additional washes to develop the colorimetric reaction. The reaction was stopped with 3M sulfuric acid 5-10 minutes after OPD addition and absorbance values were measured at a wavelength of 492nm on a spectrophotometer (Molecular Devices Emax system, Sunnyvale, CA). Absorbance values were normalized by subtracting normal wells from antigen-coated wells at the corresponding serum dilution. A sumOD value > 0.6 was required to denote a positive sample. End-point titers were defined as the reciprocal of the last adjusted serum dilution with a value ≥ 0.20.

### Serum neutralization assay

Neutralization titers were calculated by determining the dilution of serum that reduced 50% of plaques (PRNT50). We incubated a standard 100 PFU amount of SARS-CoV-2 with two-fold serial dilutions of serum samples for one hour. The virus-serum mixture was then used to inoculate Vero E6 cells for 60 minutes. Cells were overlaid with EMEM agar medium plus 1.25% Avicel, incubated for 2 days, and plaques were counted after staining with 1% crystal violet in formalin.

### Bead-based cytokine and coagulation immunoassays

Concentrations of immune mediators and fibrinogen were determined by flow cytometry using LegendPlex multiplex technology (BioLegend). Serum levels of cytokines/chemokines and plasma levels of fibrinogen were quantified using a Nonhuman Primate Inflammation 13-plex (1:4 dilution) or Human Fibrinolysis (1:10,000 dilution) panel, respectively. Samples were processed in duplicate following the kit instructions and recommendations. Following bead staining and washing, 1500-4000 bead events were collected on a FACS Canto II cytometer (BD Biosciences) using BD FACS Diva software. The raw. fcs files were analyzed with BioLegend’s cloud-based LEGENDplex™ Data Analysis Software.

### Histopathology and immunohistochemistry

Necropsy was performed on all subjects euthanized at day 5. Tissue samples of all major organs were collected for histopathologic and immunohistochemical (IHC) examination and were immersion-fixed in 10% neutral buffered formalin. Specimens were processed and embedded in paraffin and sectioned at 5 μm thickness. For IHC, specific anti-SARS immunoreactivity was detected using an anti-SARS nucleocapsid protein rabbit primary antibody at a 1:800 dilution for 60 minutes (Novusbio NB100-56683). The tissue sections were processed for IHC using the ThermoFisher Scientific Lab Vision Autostainer 360 (ThermoFisher Scientific). Secondary antibody used was biotinylated goat anti-rabbit IgG (Vector Laboratories, Burlingame, CA) at 1:200 for 30 minutes followed by Vector Streptavidin Alkaline Phosphatase at a dilution of 1:200 for 20 min (Vector Laboratories, Burlingame, CA). Slides were developed with Bio-Red (Biopath) for 7 minutes and counterstained with hematoxylin for one minute.

SARS-CoV-2 RNA *in situ* hybridization (ISH) in formalin-fixed paraffin embedded (FFPE) tissues was performed using the RNAscope 2.5 high definition (HD) RED kit (Advanced Cell Diagnostics, Newark, CA) according to the manufacturer’s instructions. 20 ZZ probe pairs targeting the genomic SARS-CoV-2 spike protein (S) gene were designed and synthesized by Advanced Cell Diagnostics (catalogue 848561). After sectioning, deparaffinization with xylene and graded ethanol washes and peroxidase blocking, the sections were heated in RNAscope target retrieval reagent buffer (Advanced Cell Diagnostics catalogue 322000) for 15 minutes and then air-dried overnight. The sections were then digested with Protease III (catalogue 322340) at 40C in the HybEZ oven (HybEZ, Advanced Cell Diagnostics catalogue 321711) for 15 minutes. Sections were exposed to ISH target probe and incubated at 40C in the HybEZ oven for 2 hours. After rinsing, the signal was amplified using the manufacturer provided pre-amplifier and amplifier conjugated to alkaline phosphatase and incubated with a red substrate-chromogen solution for 10 minutes, counterstained with hematoxylin, air-dried, and cover slipped.

## Supporting information

Supplemental Table 1_17May20

Supplemental Table 2_17May20

Supplemental Figures_17May20_1332

## Acknowledgments

The authors would like to thank the UTMB Animal Resource Center for veterinary support for surgery to implant temperature data loggers and husbandry support of laboratory animals We also thank Drs. Vineet Menachery, Shinji Makino, and Chien-Te (Kent) Tseng for their valuable insight and technical assistance with coronavirus protocols. The virus used in this publication was kindly provided by the European Virus Archive goes Global (EVAg) project that has received funding from the European Union’s Horizon 2020 research and innovation program under grant agreement No 653316. This study was supported by funds from the Department of Microbiology and Immunology, University of Texas Medical Branch at Galveston, Galveston, TX to TWG. Operations support of the Galveston National Laboratory was supported by NIAID/NIH grant UC7AI094660.

