## Supplemental Table 1_17May20 for "Establishment of an African green monkey model for COVID-19"

### Supplementary Table 1. Clinical description and outcome of African green monkeys following SARS-CoV-2 challenge

| Subject No. | Sex | Clinical illness | Clinical pathology |
|---|---|---|---|
| AGM-1 | F | Decreased appetite (d4,5). Subject survived to study endpoint (d5). | Monocytosis (d4,5); granulocytosis (d4); > 2-fold ↑ CRP (d4); > 7-fold ↑ CRP (d5) |
| AGM-2 | F | Decreased appetite (d1-5). Subject survived to study endpoint (d5). | Lymphocytopenia (d2-5); monocytopenia (d2-5); thrombocytopenia (d4); > 2-fold ↑ in ALT (d3,4); > 2-fold ↑ in GGT (d3-5); 2-fold ↑ in CRP (d3); > 3-fold ↑ in CRP (d4) |
| AGM-3 | M | None. Subject survived to study endpoint (d5). | Lymphocytopenia (d5); thrombocytopenia (d4); monocytosis (d4); granulocytosis (d4); 2-fold ↑ in CRE (d4); > 5-fold ↑ in CRP (d4) |
| AGM-4 | F | Decreased appetite (d1-10, 12, 13, 15-17). Subject survived to study endpoint (d21). | Lymphocytopenia (d2,3); granulopenia (d2,3,5), thrombocytopenia (d2); granulocytosis (d4,12,15); monocytopenia (d21); hypoglycemia (d7,9,12); > 7-fold ↑ CRP (d4); > 3-fold ↑ CRP (d5) |
| AGM-5 | M | Decreased appetite (d8,10, 17). Subject survived to study endpoint (d21). | Monocytopenia (2,4,9); granulocytopenia (d2); > 3-fold ↑ CRP (d4) |
| AGM-6 | F | Decreased appetite (d1-10, 13-17). Subject survived to study endpoint (d21). | Lymphocytopenia (d7); monocytosis (d12); > 3-fold ↑ in CRP (d4) |

Days after SARS-CoV-2 challenge are in parentheses. Lymphocytopenia, granulocytopenia, monocytopenia, and thrombocytopenia are defined by a ≥35% drop in numbers of lymphocytes, granulocytes, monocytes, and platelets, respectively. Leukocytosis, monocytosis, and granulocytosis are defined by a two-fold or greater increase in numbers of white blood cells over base line. Fever is defined as a temperature more than 2.5 °F over baseline, or at least 1.5 °F over baseline and ≥ 103.5 °F. Hypothermia is defined as a temperature ≤3.5°F below baseline. Hyperglycemia is defined as a two-fold or greater increase in levels of glucose. Hypoglycemia is defined by a a ≥25% decrease in levels of glucose. Hypoalbuminemia is defined by a ≥25% decrease in levels of albumin. Hypoproteinemia is defined by a ≥25% decrease in levels of total protein. Hypoamylasemia is defined by a ≥25% decrease in levels of serum amylase. Hypocalcemia is defined by a ≥25% decrease in levels of serum calcium. (ALT) alanine aminotransferase, (AST) aspartate aminotransferase, (ALP) alkaline phosphatase, (CRE) Creatinine, (CRP) C-reactive protein, (Hct) hematocrit, (Hgb) hemoglobin
