## Supplemental Table 2_17May20 for "Establishment of an African green monkey model for COVID-19"

**Supplemental Table 2: Gross lung lesion severity scores in AGMs infected with SARS-CoV-2**

| Subject No. | Right Upper Lobe | Right Middle Lobe | Right Lower Lobe | Left Upper Lobe | Left Middle Lobe | Left Lower Lobe |
|---|---|---|---|---|---|---|
| **AGM-1** | -- | -- | -- | * | ** | ** |
| **AGM-2** | -- | -- | *** | -- | -- | **** |
| **AGM-3** | -- | * | ** | -- | -- | * |

-- < 10%
* 25%
** 50%
*** 75%
**** > 75%
