## Supplemental Figures_17May20_1332 for "Establishment of an African green monkey model for COVID-19"

**Supplemental Figure 1: Longitudinal temperature analysis of AGMs infected with SARS-CoV-2.** Prior to challenge, AGMs were each surgically implanted with a DST micro-T small implantable thermo logger (Star-Oddi, Gardabaer, Iceland), allowing for collection of the body temperature of each animal in 15 minute increments (96 measurements/day) throughout the course of the study. AGM-1 and AGM-3 each reached temperatures 3 dpi which were noticeably above baseline temperatures (1 day prior to challenge). The “fever peak” is colored in red. Vertical dashed lines indicate the start and end of the fever peak. Horizontal dashed lines indicate the threshold temperature for classification as fever. Black arrows on the x-axis indicate time of challenge. Determination of the window of febrile temperatures was performed visually, with comparison of temperatures at all other points during the study duration (-1 dpi to 5 dpi).

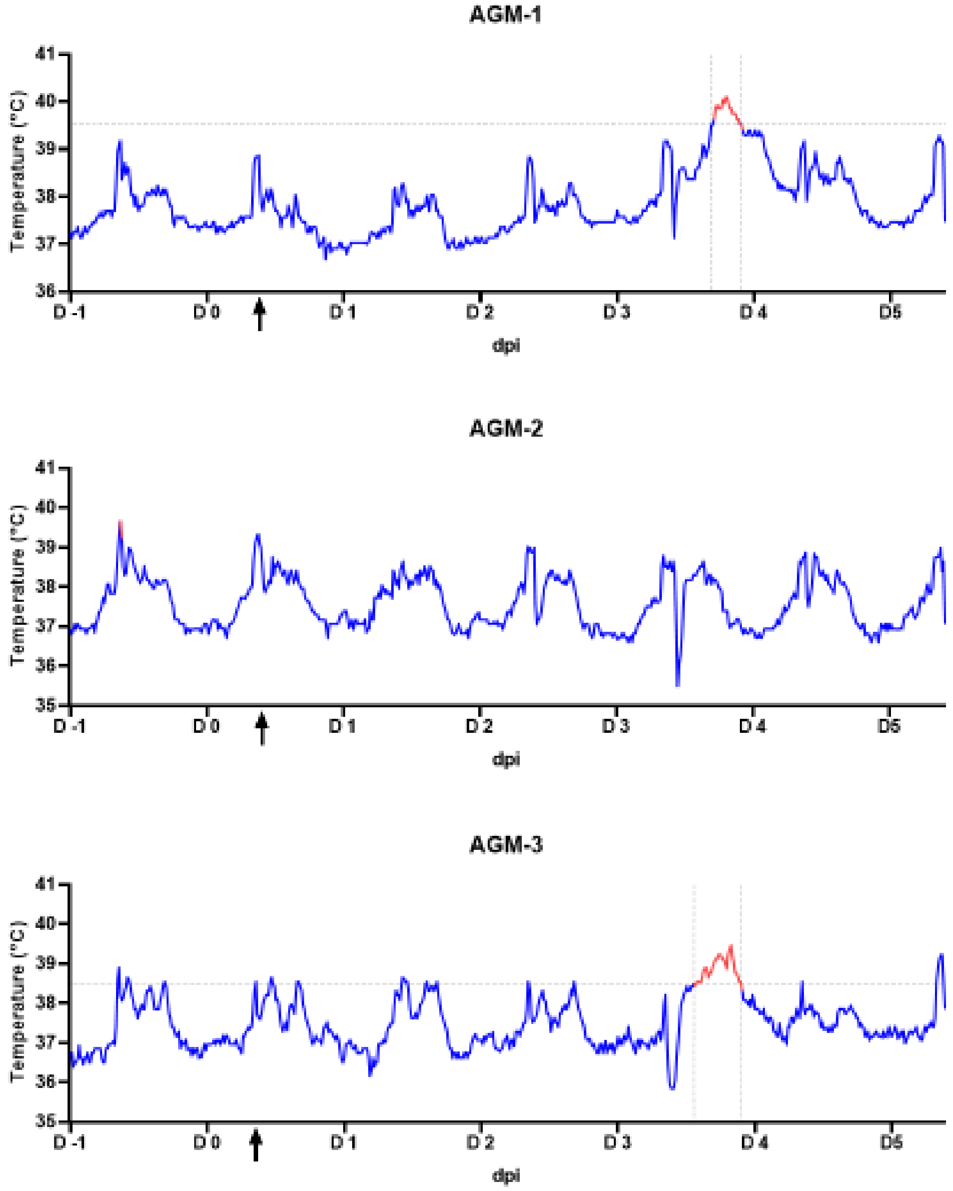

**Supplemental Figure 2: Radiographic images of time course to day 5 post infection on selected AGM**

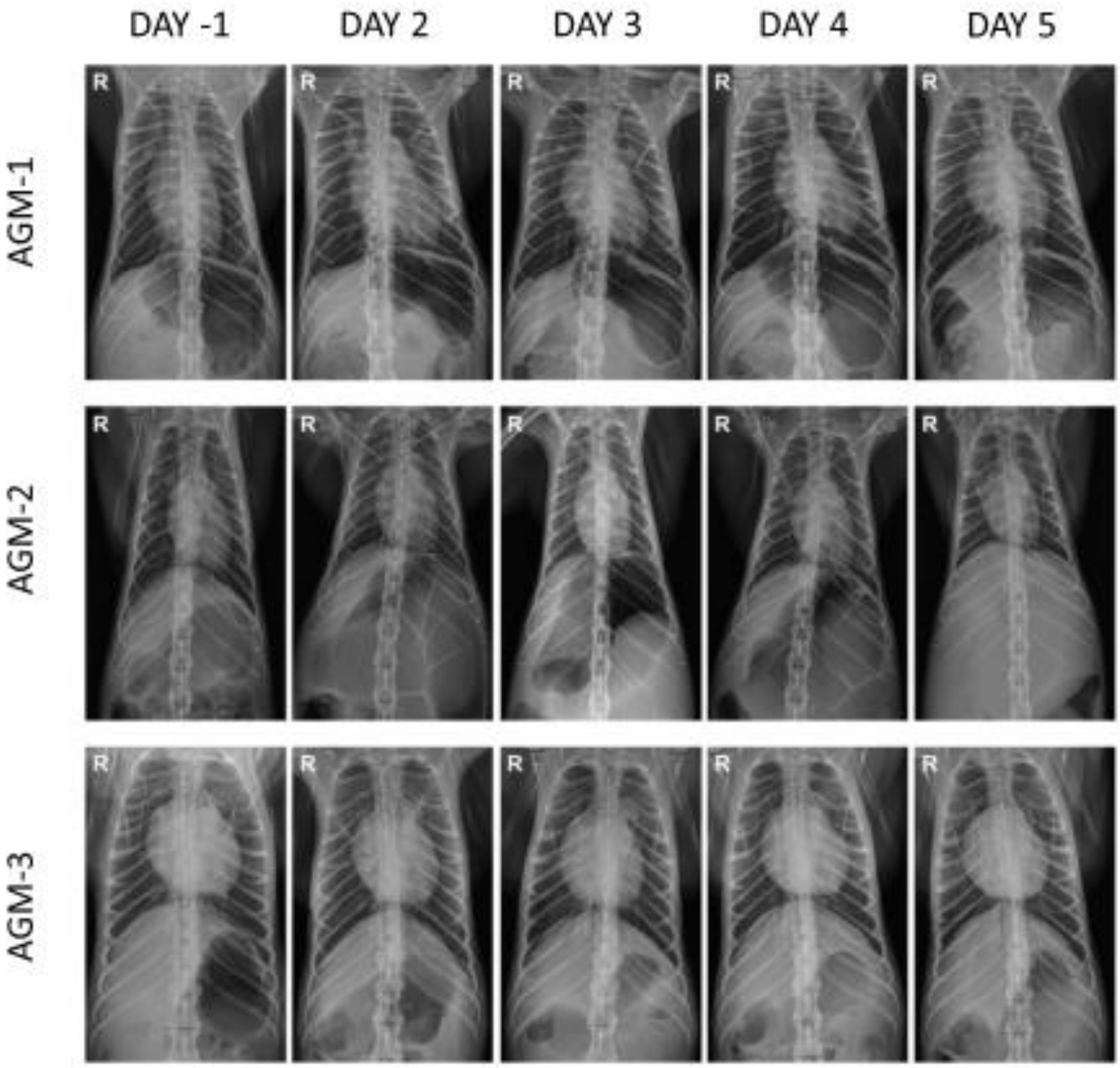
